# Why SARS-CoV-2 Omicron variant is milder? A single high-frequency mutation of structural envelope protein matters

**DOI:** 10.1101/2022.02.01.478647

**Authors:** Bingqing Xia, Yi wang, Xiaoyan pan, Xi Cheng, Hongying Ji, Xiaoli Zuo, Jia Li, Zhaobing Gao

## Abstract

SARS-CoV-2 Omicron variant is highly transmissible and extensive morbidity, which has raised concerns for antiviral therapy. In addition, the molecular basis for the attenuated pathogenicity and replication capacity of Omicron remains elusive. Here, we report for the first time that a high-frequency mutation T9I on 2-E of SARS-CoV-2 variant Omicron forms a non-selective ion channel with abolished calcium permeability and reduced acid sensitivity compared to the WT channel. In addition, T9I caused less cell death and a weaker cytokine production. The channel property changes might be responsible for the Omicron variant releases less efficiently and induces a comparatively lower level of cell damage in the infected cells. Our study gives valuable insights into key features of the Omicron variant, further supporting 2-E is a promising drug target against SARS-CoV-2 and providing critical information for the COVID-19 treatment.

## Mian Text

Since November 2021, a new SARS-CoV-2 variant (B.1.1.529) emerged in South Africa and was designated as the fifth variant of concern (VOC), named as Omicron ^1, 2^. Omicron variant is highly transmissible and extensive morbidity, which has raised concerns for antiviral therapy. Omicron has largest number (>30) of substitutions, deletions, or insertions with mutations frequency higher than 1% in the spike protein (S). Mutation studies in the spike RBD domain have provided a plausible explanation for altered transmissibility and antibody resistance ^3–5^. Recent reports revealed Omicron variant induces a longer cycle for virus shed, attenuates replication capacity, and produces substantially attenuated lung pathology, indicating that the pathogenic ability of Omicron variant is indeed milder ^6–8^. However, the molecular basis for the attenuated pathogenicity and replication capacity remains elusive. The envelope protein of SARS-CoV-2 (2-E) forms a homo-pentameric cation channel that is important for virus pathogenicity^9, 10^. It has been demonstrated that 2-E channel alone is sufficient to induce cell death, provoke cytokine storm and even cause acute respiratory distress syndrome (ARDS)-like damages *in vivo*. Its inhibitor exhibits excellent antiviral activity *in vivo^9^.* Therefore, we focus on the 2-E mutations of Omicron variant.

Three high quality E protein sequences of Omicron variants (B.1.1.529, BA1and BA2) in the CNCB database as of January 18th, 2022 (https://ngdc.cncb.ac.cn/ncov/) were comprehensively analyzed and a mutation 2-E^T9I^ (T9I) caught our attention. Compared with the original Wuhan reference strain, among more than one hundred identified 2-E mutations in Omicron variant, T9I shows a much higher frequency than the rest mutations. In addition, the mutation probability of T9I in all statistical samples is higher than 99.5%. For the most common Omicron strain B.1.1.529, the mutation probability is 100% (Fig.1a, b). According to the solved NMR structure, T9 locates at the top of the transmembrane domain (TMD) of 2-E proteins ^10^. The influence of T9I has not been studied yet completely.

First, we asked whether T9I still retains channel activity. We purified T9I protein and reconstituted the protein on Planar Lipid Bilayer (BLM) as previously described ^9^(Supplementary Fig.S1). The observed T9I-induced typical single-channel currents supported that T9I was able to form ion channels also as wild-type (WT) 2-E proteins ^9^. However, the reversal potential of T9I channels was shifted to left, from 57 mV to 3 mV, under asymmetric KCl solutions. Similar reversal potential shifting was also detected in asymmetric NaCl solutions (Fig.1c.d, Supplementary Fig.S2). A reversal potential close to 0 mV suggests that T9I may have lost its selectivity to cations. To determine whether T9I can permeate chloride, the channel activity of T9I was further examined in asymmetric 50:500 mM choline chloride solutions. The quaternary ammonium choline is not permeable to most cation channels and thus is always used to detect anion permeability. As expected, outward currents were indeed observed when the holding voltage potentials were higher than the theoretical reversal potential for chloride (Erev> 70 mV). Besides, WT channels are permeable to Ca^2+^. As shown in Figure S2c, for WT channels, the frequent and continuous inward potassium currents in the asymmetric 50:500 mM (trans: cis) K^+^ solutions indicated that the 2-E channels were incorporated into the membranes. The membrane potential was then changed to b+65 mV (K^+^ reverse potential) to eliminate the K^+^ currents. Intriguingly, outward step-like signals appeared when 20 mM (final concentration) Ca^2+^ was added to the *trans* side. When the Ca^2+^ concentration increased to 20 mM, the outward currents increased. In contrast, under the identical conditions, T9I channels did not induce detectable calcium currents (Supplementary Fig.S2c). These results revealed that T9I owns a different ion selectivity and permeability from those of WT.

WT channels are pH sensitive ^9^. The pH influences on T9I channels were evaluated on a same channel using titration strategy. After the channels incorporated into the membranes, HCl was titrated into either *cis* or *trans* side to alter the pH to 4 from 6. Consistent to our previous results, reduction of pH, either in *cis* or *trans* side, induced significant increase of amplitude and open probability of WT channels. In contrast, pH reduction in *cis* side failed to increase the channel activity of T9I channels. Although we didn’t have direct evidences to explain how the threonine (T9) site is sensitive to pH changes, the structure of 2-E channels supported the residues (Glu7 and Glu8) around T9 were sensitive to pH change and their side chain carboxyl could be deprotonated at neutral pH and protonated at acidic pH ^10^. It is reported that SARS-CoV-2 traffics to the lysosomes for egress by lysosome deacidification, instead of using the conventional biosynthetic secretory pathway ^11^. The E channels of coronavirus are localized to intracellular organelles, including lysosome and Golgi apparatus ^11, 12^. The cellular localization and expression of WT and T9I were assessed using immunocytochemistry and confocal microscopy. We found that both WT and T9I are co-localized with the lysosomal marker LAMP (Fig.1g, supplementary Fig.S3). Given the localization on lysosome, the influences of WT and T9I on the luminal pH of lysosome were compared. Flow cytometric was used to analyze pH probe (pHluorin) ratio. This pH probe consists of green fluorescent protein (GFP) pHluorin molecule fused with lysosome-resident protein CD63. To generate a pH calibration curve, un-transfected cells were subjected to treatment with buffers ranging from pH 5.5 to 7.5 in the presence of the ionophores monensin and nigericin prior to flow analysis (Fig.1h, i). Expression of WT channels robustly neutralized the lysosome pH, whereas T9I channels exhibited less influences in luminal pH, which was consistent with our electrophysiological results (Fig.1j). Therefore, the capability of deacidification of WT channels in lysosomes was significantly impeded by the mutation T9I. Based on above, whether the weakened capability of deacidification hampers virus releasing was examined. We transfected the Vero E6 cells with WT or T9I plasmids at day 1, then the cells were infected with SARS-CoV-2 virus at day 2. As determined by qRT-PCR, pre-expression of mutant T9I significantly attenuated viral loadings compared with WT (Fig.1k). It was logic to deduce that some exogenous T9I subunits would co-assemble with the WT subunits originated from the virus and form heteromultimers with impaired pH sensitivity, resulting in less deacidification of lysosomes and less SARS-CoV-2 virus releasing.

Finally, the channel activity of the 2-E protein is a determinant of virulence. We used Annexin V/PI staining and CCK8 assays to evaluate the cell lethality of T9I in Vero E6 cells. We found that although the expression level of T9I was far exceeding the expression level of WT (Fig. 1 l), it caused similar cell death as WT did (Fig. 1 m, supplementary Fig.S4a). Similarly, in the mouse macrophage cell line (RAW264.7), a very highegreater expression of T9I induced a comparable increase of inflammatory cytokines as WT did (Fig. 1n, Supplementary Fig.S4b). The correlation between the cell lethality and the expression level was further compared. As shown in Figure 1o, in comparison with WT, the killing cells capability of T9I was dropped around 150 times as well as the cytokines production. Notably, according to another latest study, the 2-E expression in Omicron variant was decreased approximately 3-fold, which may further suppress the virulence of Omicron variants ^6^. How much the weakened capabilities of killing cells and inducing cytokine production confer the milder clinical symptoms of Omicron warrant further investigation.

**Fig.1.**
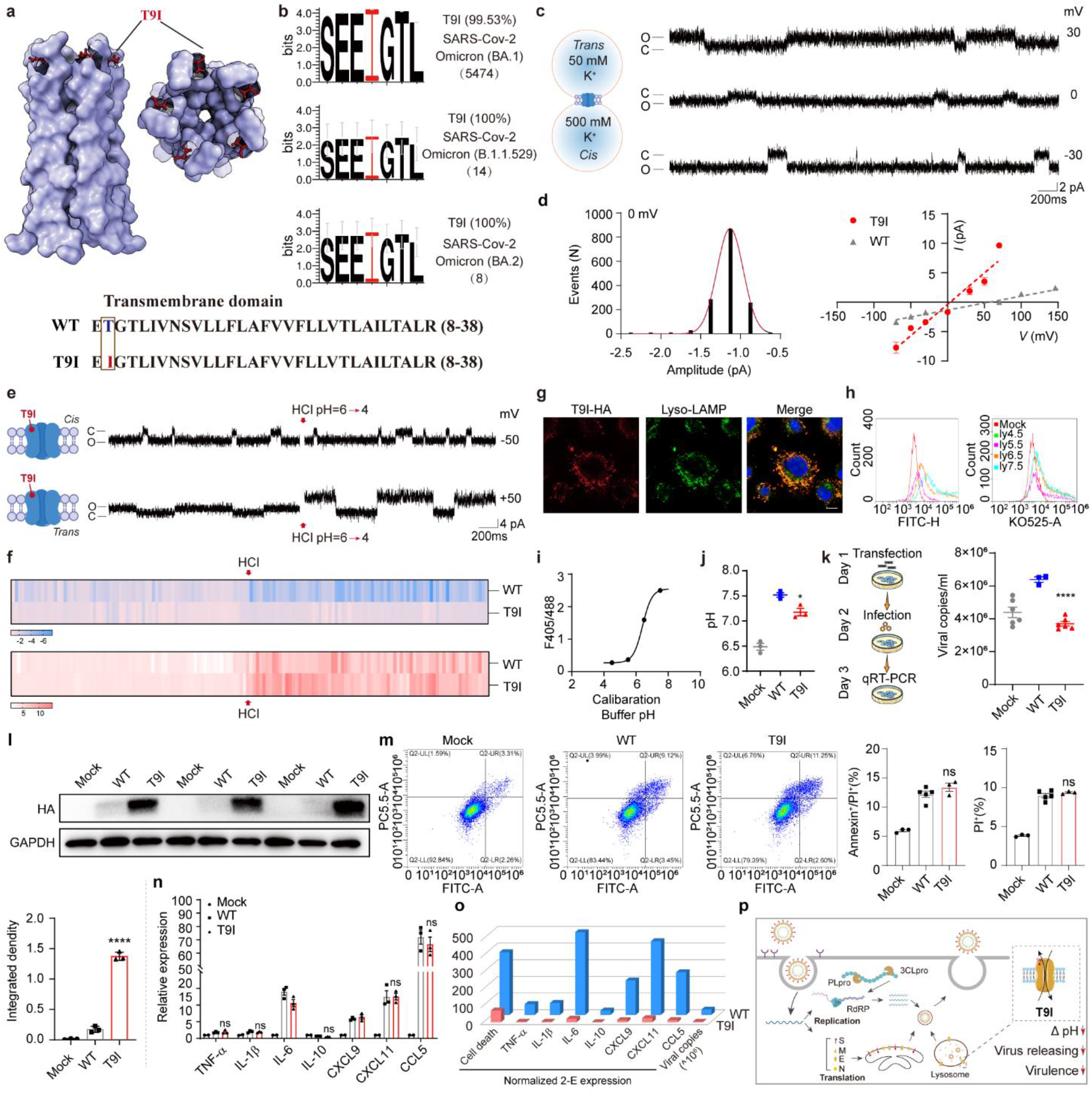
The envelope protein mutation T9I of SARS-CoV-2 Omicron variant shows less pathogenicity and weakens virus production. **a,** Structure of the 2-E TM domain ^10^. Illustration of the amino-acid sequence of SARS-CoV-2 E (8-38) and the indicated mutants. Threonine residues (T) are marked as blue and the T>I mutants are marked as red. **b,** 2-E protein amino-acid sequence logo of SARS-CoV-2 Omicron variant. The I9 residues are marked as red. 2-E protein sequences were downloaded from China National Center for Bioinformation (CNCB) using the “Lineage Browse” option. Logos were generated using the WebLogo 3.7.4 online tool. **c,** Single-channel current recording of T9I after reconstitution in lipid bilayers with PC/PS = 3:2 lipids at indicated potentials and solutions (Left). **d,** Left, all-point current histograms for the trace in **c** (0 mV) (“C” means Closed; “O” means Open). Right, *I-V* curves of T9I in the solutions of **c** (n ≥ 3). **e**, Single-channel current recording in 500 :500 mM KCl solutions via proton titration result in different pH (*cis* or *trans*). Red arrow represents titration of HCl. **f,** The heat map shows the transition of the currents from panel **e** under proton titration. **g,** Cell immunofluorescence of T9I (Red) and lysosome marker LAMP (Green). T9I is co-localized with lysosome. **h-j,** Lyso-pH indicator was used to assess the pH of the lysosome via flow cytometry. Flow cytometry pattern of the cells in buffers of known pH and contained ionophores to equilibrate the extracellular and lysosome luminal pH (**h**). Calibration curves were generated from data like those illustrated in h (**i**). **j,** The pH value of Vero E6 cells lysosome transfected with WT, T9I or Mock. **k.** Flow chart of the experiment (Left), the virus copies in supernatant determined by qRT-PCR assay (Right). MOI of SARS-CoV-2 for infection was 0.01. **l,** Protein expression levels of WT and T9I in Vero E6 cells after transfection (Up). The histogram shows the quantization of the result (Bottom). **m,** Flow cytometry pattern of Vero E6 cells transfected with WT or T9I plasmids. Right, flow cytometry analysis of Propidium iodide (PI) and Annexin V stained Vero E6 cells. **n,** Serum cytokine levels at 24 h after transfection with WT, T9I or Mock detected by RT-qPCR. **o,** The relative activity of WT, T9I in pH regulation, cell virulence, channel conductance and viral copies normalized to the expression levels. **p,** Life cycle pattern diagram of SARS-CoV-2 Omicron variant. T9I on envelope protein (2-E) of SARS-CoV-2 variant Omicron forms a non-selective ion channel with weakened pathogenicity and virus production. Experiments were independently performed thrice and similar results were obtained. One set of reprehensive data is shown here (n ≥ 3). \**p* < 0.05; \*\**p* < 0.01; \*\*\**p* < 0.001; unpaired Student’s t test. All error bars are SEM.

Overall, as SARS-CoV-2 infections are progressing, there are chances that virus accumulates new mutations. Among existed antiviral targets, different from spike protein, RNA-dependent RNA polymerase (Rdrp) and 3CL pro, which mediate virus entry, protein cleavage and viral replication, respectively, the envelope protein (2-E) majorly participating in the virus releasing ^13–15^. Here we report for the first time that a high-frequency 2-E mutation T9I of SARS-CoV-2 variant Omicron forms a non-selective ion channel with abolished calcium permeability and reduced acid sensitivity compared to WT channels. In addition, T9I caused less cell death and a weaker cytokine production (Fig.1p). The channel property changes might be responsible for the Omicron variant releases less efficiently and induces a comparatively lower level of cell damage in the infected cells. According to 2-E NMR structures, although T9I is not located in the pore domain of the channel, it may alter the channel configuration via interacting with the amino acids at positions L12 and T11 (Supplementary Fig.S5). Our study provides valuable insights into key features of the Omicron variant and critical information for the COVID-19 treatment.

## Acknowledgement

We are grateful to the National Science Fund of Distinguished Young Scholars (81825021), Fund of Youth Innovation Promotion Association (2019285), the National Natural Science Foundation of China (81773707, 92169202), the National Key Research and Development Program of China (2020YFC0842000), the National Key Laboratory Program of China (LG202101-01-04), Fund of National Science and Technology Major Project (2018ZX09711002-002-006) and the Hubei Science and Technology Project (2020FCA003) for financial support.

## Author contributions

Z. G., and B. X. conceived designed the project. Z. G., J. L., and B. X designed the experiments; Y. W. and B. X. performed the electrophysiological recordings; B. X. carried out the cell-based assays; P. X. carried out the virus assays in vitro; Y. W purified the proteins; H.Y. analyzed the sequence; all authors analyzed and discussed the data. Z. G., B. X. wrote the manuscript. All authors read and approved the manuscript.

## Method

### Plasmids and mutagenesis

Wild type SARS-CoV-2-E sequences were synthesized by the Beijing Genomics Institute (BGI, China). The mutants 2-E^T9I^ sequence was generated by site-directed mutagenesis and confirmed by sequencing (BGI, China). Vector pET28a was used for protein purification; vector pcDNA5 and pcDNA3.1 were used for cell survival assay and cell imaging.

### Cell culture and treatment

Vero E6 and Raw 264.7 cells were grown in 90% DMEM basal medium (Gibco, USA) supplemented with 10% fetal bovine serum (Gibco, USA) at 37°C under 5% CO_2_.

### Planar lipid bilayers recording

According to our previous methods, the purified protein 2-E^WT^ or 2-E^T9I^ were incorporated into lipid bilayers to test their functionality. The lipid bilayers were prepared by PC: PS= 3: 2 (Avanti Polar Lipids, USA). Proteins were added in *cis* side and the solution was 50:500 mM (*trans*: *cis*) KCl or others, all solutions were buffered by 5mM HEPES, pH 6.35. Membrane currents were recorded by headstage connected with amplifier BC-535 (Warner Instruments, USA), filtered at 1–2 kHz and digitized by Axon Digidata 1440A (Molecular Devices, USA). The recording frequency was 10 kHz. The data were processed using pClamp 10.2 software (Molecular Devices, US). The single-channel conductance was determined by fitting to Gaussian functions (bin width = 0.25 pA). Opening time less than 0.5–1.5 ms was ignored to avoid noise interference. The equilibrium potential was calculated using Goldman–Hodgkin–Katz flux equation:

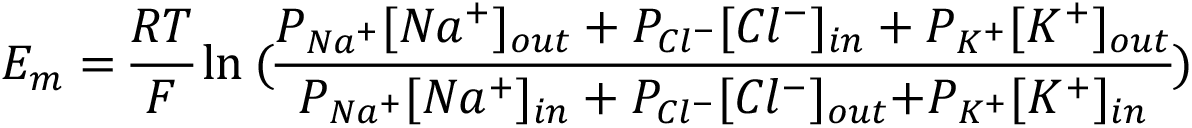

### Cell viability and cytotoxicity assay

Cell viability was measured by the CCK-8 kit (40203ES60, Yeason, China) according to the manufacturer’s instructions. The absorbance was measure at 450 nm with Thermo Scientific Microplate Reader (Thermo Fisher Scientific Inc., USA).

### Western blotting and antibodies

Proteins were resolved in 12% SDS-PAGE, transferred to PVDF membranes (GE, USA). The PVDF membranes were incubated with primary antibodies against HA-Tag (C29F4) (3724, CST, USA) or GAPDH (30201ES20, Yeasen, China), then incubated with second antibodies against IgG (Yeasen, China).

### Flow cytometry of cell death

After treatment, cells were detached, collected by centrifugation and resuspended in 1× binding buffer containing 100 μg/mL propidium iodide (PI) and Annexin V-FITC (1:20, V13242, Life, USA) and incubated at room temperature for 15 min in the dark. Subsequently, samples were added 400 μL of 1× binding buffer and kept on ice. Cells were analyzed by flow cytometry using BD FACSCALIBUR 4 (BD COMPANY, USA). The percentages of differently labeled cells were calculated by FlowJo 7.6.

### Immunofluorescence microscopy

Vero E6 cells were seeded as 1×10^4^ cells per well in coverslips on chamber slides (Thermo Fisher Scientific Inc., USA) overnight. Cells were transfected with WT or T9I with lysosome marker (LAMP1,). After transfection for 24 h, cells were washed with phosphate-buffered saline (PBS) and fixed in 4% paraformaldehyde for 10 min at room temperature. Then the cells were permeabilized in 0.5% Triton X-100 in PBS for 3 min. The coverslips were washed twice with PBS and incubated with primary antibody in 1% bovine serum albumin (BSA) (36101ES25, Yeasen, China) for 2 h at room temperature. Anti-HA (HT301-01, Trans, China) and rabbit anti-LAMP (55273-1-AP, Proteintech, USA) were used to incubate the cells at a 1:1,000 dilution. Then, second antibodies FITC-AffiniPure Goat Anti-Rabbit IgG (33107ES60, Yeasen, China) and Cy3-AffiniPure Goat Anti-Mouse IgG (33208ES60, Yeasen, China)were used to incubate the cells at a 1:1,000 dilution. Images were captured using the Leica TCS-SP8 STED system with 100× oil objective.

### Virus loading and quantitative real-time PCR (qRT-PCR) analysis

SARS-CoV-2 (nCoV-2019BetaCoV/Wuhan/WIV04/2019) was preserved at Wuhan institute of virology, Chinese Academy of Sciences. It was propagated and titrated with Vero E6 cells, and its associated operations were performed in a biosafety level 3 (BSL-3) facility. Cell supernatants RNA was isolated with MiniBEST Viral RNA/DNA Extraction Kit (Takara, Japan) as described in the instruction, and cDNA was transcribed with PrimeScript™ RT reagent Kit with gDNA Eraser (Takara, Japan). In detail, 50 μL supernatant were collected for RNA isolation with a MiniBEST Viral RNA/DNA Extraction Kit Ver.5.0 (Takara, AK41820A), and the total RNA was eluted with 30 μL RNase-free water. cDNA was transcribed from 3 μL total RNA in 20 μL reaction system with PrimeScriptTM RT reagent Kit with gDNA Eraser (Takara, AK71648A). Viral copies were quantified from 1 μL template viral cDNA by a standard curve method on ABI 7500 (Takara TB Green Premix Ex Taq II, AK81975A) with a pair of primers targeting S gene. The forward primer (5’-3’) is: CAATGGTTTAACAGGCACAGG; the reverse primer (5’-3’) is: CTCAAGTGTCTGTGGATCACG.

### Determination of lysosome pH

Vero cells were transfected with Lyso-pHluorin (addgene,70113) and 2-E plasmid. After transfection, cells were trypsinzed and washed with Live Cell Imaging Solution (LCIS). Then, replaced LCIS with cellular pH calibration Buffers containing 10 μM Valinomycin and Nigericin (Intracellular pH Calibration Buffer Kit, P35379, Themo, USA) and incubate at 37°C for at least 5 minutes. At last, cells were analyzed via Flow cytometry with excited at 405 nm and 488 nm, and the emission signals were collected with detection filters at 500 to 550 nm and 515 to 545 nm, respectively. Flow cytometric data were collected and quantified using CytoExpert 2.4 software.

### Homology modeling

The solid-state NMR structure of SARS-CoV-2 envelope protein (PDB code: 7k3g) [(2020) Nat Struct Mol Biol 27: 1202-1208] was used to as the template to build homology models of the mutant envelope protein. This solid-state data yield to ten lowest-energy structures. These ten structures were employed to construct ten homology models of the mutant envelope protein using Modeller [A. Sali, T.L. Blundell, Comparative protein modelling by satisfaction of spatial restraints, J Mol Biol 234 (1993) 779-815. 10.1006/jmbi.1993.1626.]. The models with the lowest root mean square deviations from their template structures were selected.

### Statistics

All measurements were derived from distinct samples. Statistics were performed in GraphPad Prism. Statistical significance was determined by one-way ANOVA and two-way ANOVA followed by pairwise Student’s t-test with Tukey’s or Sidak’s correction. Two-tailed unpaired Student’s t test was performed if only two conditions were compared. Data in the text is presented as Mean±SEM. Adjusted P values are reported in the figures.

## Supplementary figure

**Supplementary information, Fig. S1.**
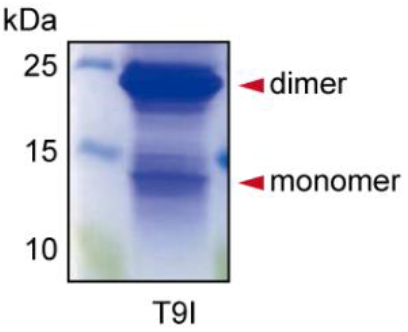
Purification of T9I proteins. Purification of full-length 2-E^T9I^ protein with Ni-NTA affinity chromatography. 15% SDS-PAGE gel with coomassie blue staining.

**Supplementary information, Fig. S2.**
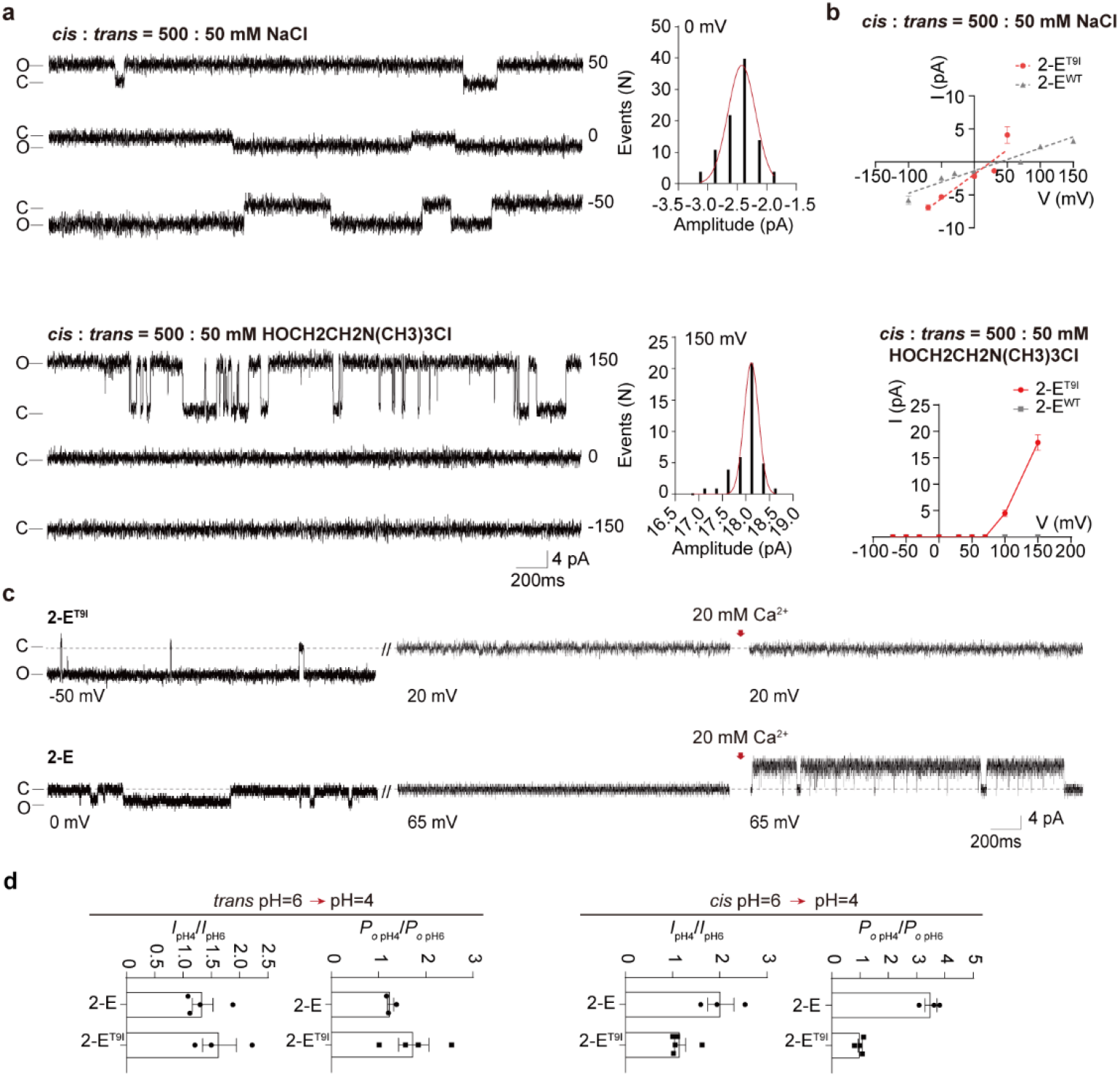
Electrophysiological property of T9I. **a.** Single-channel current recording of T9I after reconstitution in lipid bilayers with PC/PS = 3:2 lipids at indicated potentials and solutions. Protein (5–50 ng/mL) was added to the cis side. All-point current histograms for the left trace (0, 150mV) (“C” means Closed; “O” means Open). **b.** *I-V* curves of T9I in indicated solutions (n ≥ 3). **c.** Pure Ca^2+^ currents recording in 50 :500 mM KCl solutions (*trans*: *cis*) via calcium titration as indicated concentrations, red arrow represents titration of CaCl2. **d.** Amplitude (*I*) and open probability (*Po*) of channel current at pH 4 were normalized to channel current at pH 6 of both sides (n ≥ 3).

**Supplementary information, Fig. S3.**
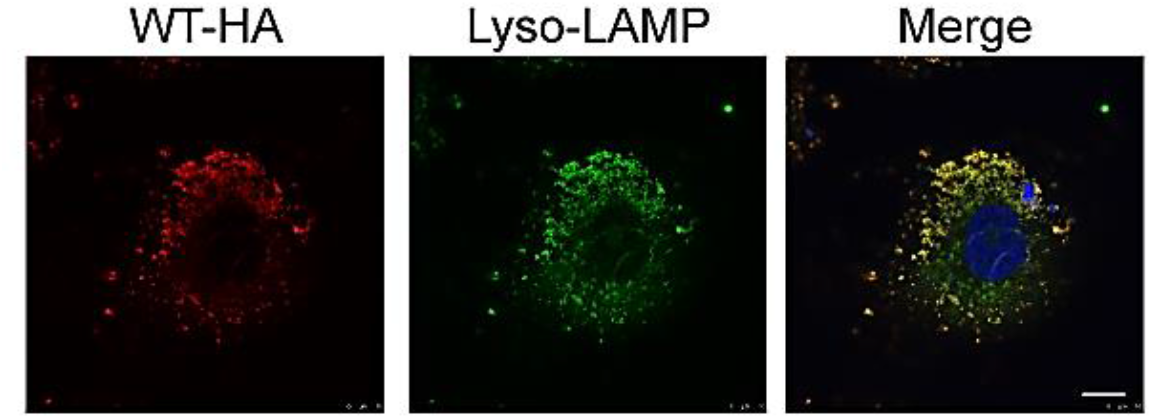
Wild type envelope protein (E) is co-localized with lysosome. Cell immunofluorescence of WT (Red) and lysosome marker (Green). Scale bar,10 μM.

**Supplementary information, Fig. S4.**
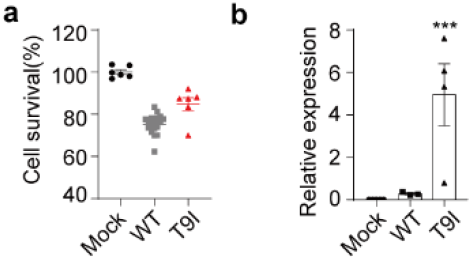
Cell viability of Vero E6 cells after transfection with 2-E plasmids and expression level of after transfection with WT andT9I. **a.** Cell viability of Vero E6 cells after transfection with WT, T9I or Vector (Mock) at 24 h. **b.** Expression level of 2-E after transfection with WT, T9I or Mock in Raw 264.7 cells detected by RT-qPCR.

**Supplementary information, Fig. S5.**
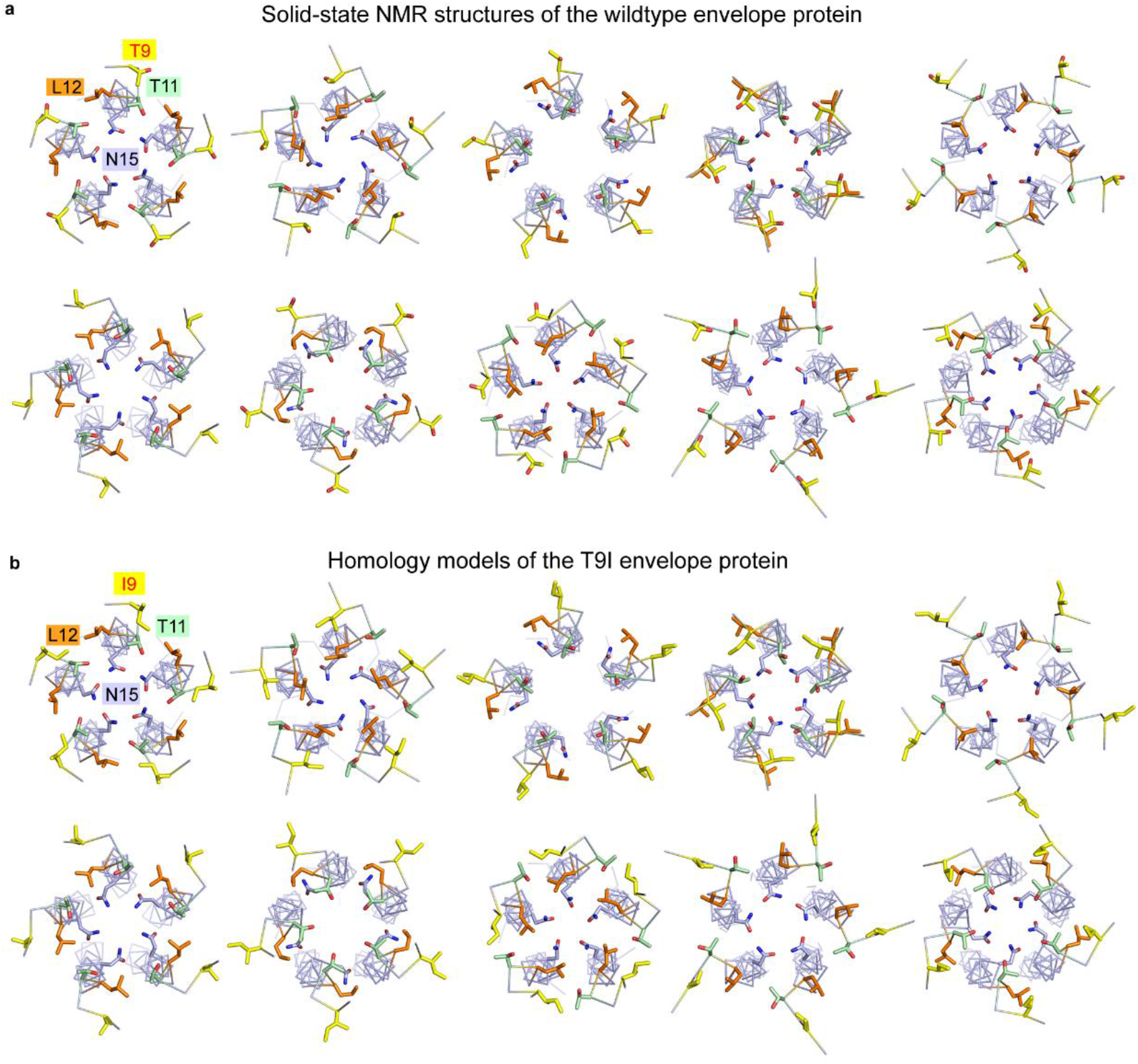
Structure of WT and T9I. **a**, Top view of ten lowest-energy solid-state NMR structures of wildtype SARS-CoV-2 E protein (PDB code: 7k3g). Three N-terminal residues T9, T11, L12 and a pore-lining residue N15 are shown as sticks (T9 in *yellow*, T11 in *green*, L12 in *orange*, N15 in *light blue*). The protein is shown as ribbon. **b,** Top view of ten homology models of2-E^T9I^, which built based on the solid-state NMR structures of wildtype SARS-CoV-2 envelope protein (PDB code: 7k3g). Three N-terminal residues I9, T11, L12 and a pore-lining residue N15 are shown as sticks (I9 in *yellow*, T11 in *green*, L12 in *orange*, N15 in *light blue*).

